# Community composition of coral-associated Symbiodiniaceae is driven by fine-scale environmental gradients

**DOI:** 10.1101/2021.11.10.468165

**Authors:** Mariana Rocha de Souza, Carlo Caruso, Lupita Ruiz-Jones, Crawford Dury, Ruth Gates, Robert J. Toonen

## Abstract

The survival of reef-building corals is dependent upon a symbiosis between the coral and the community of Symbiodiniaceae. *Montipora capitata*, one of the main reef building coral species in Hawaiʻi, is known to host a diversity of symbionts, but it remains unclear how they change spatially and whether environmental factors drive those changes. Here, we surveyed the Symbiodiniaceae community in 600 *M. capitata* colonies from 30 sites across Kāneʻohe Bay and tested for host specificity and environmental gradients driving spatial patterns of algal symbiont distribution. We found that the Symbiodiniaceae community differed markedly across sites, with *M. capitata* in the most open-ocean (northern) site hosting few or none of the genus *Durusdinium*, whereas individuals at other sites had a mix of *Durusdinium* and *Cladocopium*. Our study shows that the algal symbiont community composition responds to fine-scale differences in environmental gradients; depth and temperature variability were the most significant predictor of Symbiodiniaceae community, although environmental factors measured in the study explained only about 20% of observed variation. Identifying and mapping Symbiodiniaceae community distribution at multiple scales is an important step in advancing our understanding of algal symbiont diversity, distribution and evolution, and the potential responses of corals to future environmental change.

## 1. Introduction

Coral reefs are among the most biologically diverse and productive ecosystems on Earth and provide valuable ecosystem services as sources of tourism, coastal protection, natural products, and nutrition (Hicks, Graham, & Cinner, 2013; Moberg & Folke, 1999; Woodhead, Hicks, Norström, Williams, & Graham, 2019). The symbiotic interaction with an exceptionally diverse dinoflagellate (family Symbiodiniaceae) is inherently linked to the health and success of reef-building corals, because they provide a large proportion of the coral energy requirement (Muscatine & Porter, 1977). There are eleven genera (Yorifuji et al., 2021) of Symbiodiniaceae, each with different physiological characteristics that impact the nutrient provisioning and thermal tolerance of the coral host (Berkelmans & van Oppen, 2006; Grégoire, Schmacka, Coffroth, & Karsten, 2017; Hoadley et al., 2015; Howells et al., 2012; McIlroy et al., 2016; Quigley, Baker, Coffroth, Willis, & van Oppen, 2018; Robison & Warner, 2006; Stat & Gates, 2011). *Cladocopium* (previously *Symbiodinium* clade C) and *Durusdinium* (previously clade D) are the two genera most commonly hosted by corals in the Pacific ( LaJeunesse, 2005). *Cladocopium* is a generalist symbiont and also the most speciose genus (Baker, 2003; LaJeunesse et al., 2018) while *Durusdinium* is usually found in shallow corals exposed to high light or sea surface temperature or areas with high temperature variability (Oliver & Palumbi, 2011) and is associated with increased resilience to thermal stress (Berkelmans & van Oppen, 2006; LaJeunesse et al., 2018, 2010; Oliver & Palumbi, 2011; Rowan, 2004; Stat & Gates, 2011).

Thermal stress, mostly caused by climate change, is currently the main threat affecting corals worldwide (Heron, Maynard, van Hooidonk, & Eakin, 2016; Hoegh-Guldberg et al., 2007; Hughes, 2003; Hughes et al., 2017; Sully, Burkepile, Donovan, Hodgson, & van Woesik, 2019). Sea temperatures in many tropical regions have increased by almost 1°C over the past 100 years and are currently increasing at ~1-2°C per century (Lough, Anderson, & Hughes, 2018; Peñaflor, Skirving, Strong, Heron, & David, 2009; Sully et al., 2019; IPCC 2021). Temperature stress disrupts coral-dinoflagellate symbiosis, leading to algal symbiont loss and consequent paling, a phenomenon known as coral bleaching (Brown, 1997; Douglas, 2003). Mass coral bleaching events are increasing in frequency and duration, resulting in significant losses of live coral in many parts of the world (Donner, Skirving, Little, Oppenheimer, & Hoegh-Guldberg, 2005; Hoegh-Guldberg, 1999; Oliver, Berkelmans, & Eakin, 2018; Sully et al., 2019). Coral susceptibility to heat stress and bleaching is dependent on a wide range of factors, including the algal symbiont community they host (Berkelmans & van Oppen, 2006; Douglas, 2003; Hoegh-Guldberg et al., 2007; Rowan, 2004; Warner, LaJeunesse, Robison, & Thur, 2006). Bleaching may also represent an opportunity for corals to rapidly change their current algal symbiont community composition to more resilient types (Buddemeier & Fautin, 1993; Fautin & Buddemeier, 2004). However, this Adaptive Bleaching Hypothesis remains controversial because many coral taxa are algal symbiotic specialists, hosting a single algal symbiont taxon and although corals can sometimes change their Symbiodiniaceae symbionts, many also recover to the same algal symbiont community they had previous to bleaching (Baird, Cumbo, Leggat, & Rodriguez-Lanetty, 2007; Goulet, 2006, 2007; Goulet & Coffroth, 2003; Hoegh-Guldberg, Jones, Ward, & Loh, 2002). Furthermore, hosting the stress tolerant *Durusdinium* often comes at an energetic cost, as it decreases the growth and metabolite exchange rate of the host (Baker, Andras, Jordán-Garza, & Fogel, 2013; Cunning, Silverstein, & Baker, 2018; Jones & Berkelmans, 2010; Jones & Berkelmans, 2011; Matthews et al., 2018; Sproles et al., 2020, 2019).

Symbiodiniaceae assemblage structure in corals tend to be shaped by many factors, including the host species (Baker, 2003; Finney et al., 2010), large scale factors like geography (LaJeunesse et al., 2004; Tonk, Sampayo, Weeks, Magno-Canto, & Hoegh-Guldberg, 2013), and local scale factors like depth (Bongaerts, Ridgway, Sampayo, & Hoegh-Guldberg, 2010; Warner et al., 2006), habitat (LaJeunesse et al., 2010), and environmental factors such as light (Rowan, Knowlton, Baker, & Jara, 1997) and temperature (LaJeunesse et al., 2010; Tonk et al., 2013). Here, we investigated the local-scale environmental drivers of Symbiodiniaceae assemblage structure in a common reef-building coral *Montipora capitata*. *Montipora capitata* is one of the most abundant corals in Kāne‘ohe Bay and may harbor a community of *Cladocopium* (C), *Durusdinium* (D) or both symbiont genera (Cunning, Ritson-Williams, & Gates, 2016; Dilworth, Caruso, Kahkejian, Baker, & Drury, 2021; Innis, Cunning, Ritson-Williams, Wall, & Gates, 2018). Previous studies of the species in Hawaiʻi have reported associations with C31 (Cunning et al., 2016; Stat et al., 2011, 2013), C17, C21 (Stat et al., 2011, 2013) and *Durusdinium glynnii* (formerly, *Symbiodinium glynii* [ITS2 Type D1 (Stat et al., 2011, 2013)], and D 4-6 (Cunning et al., 2016).

We used high-throughput sequencing of the internal transcribed spacer region (ITS2) between RNA genes to identify the assemblage and the local-scale environmental drivers of Symbiodiniaceae of 600 colonies of the common reef-building coral *Montipora capitata* collected from across 30 sites in Kāneʻohe Bay. Kāneʻohe Bay is the largest sheltered body of water in the Main Hawaiian Islands, and reefs across the bay experience different daily ranges in temperature, pH, irradiance, and oxygen concentration as well as oceanic influence, water turnover time, and freshwater and nutrient input (Bahr, Jokiel, & Toonen, 2015). We collected data on temperature, sedimentation, and depth at each site to determine the predictive value of these environmental drivers on the observed Symbiodiniaceae community. These spatial differences in environmental variability create a great study system to explore how environment gradients in different parts of the bay can shape the Symbiodiniaceae community.

## 2. Material and Methods

### 2.1 SITE SELECTION AND TAGGING

*Montipora capitata* colonies were tagged and collected under Special Activity Permit 2018-03 to HIMB from Hawai‘i Department of Aquatic Resources. Details of the sampling design and block delimitation have been described in (Caruso et al., 2021). In summary we performed stratified random sampling of sites by first dividing the bay into five blocks, based on the water flow regimes and water residence time (Lowe, Falter, Monismith, & Atkinson, 2009b, 2009a). Block 1 has the longest water residency, with > 30 days on average, blocks 2 and 3 have a residency of 10-20 days, whereas blocks 4 and 5 had typical residency times less than 1 day. We generated 20 random points in each block. The points in each block were visited in random order and the first six to have at least twenty *Montipora capitata* of 10 cm or more in diameter were retained as sampling sites, for a total of 30 sites across the 5 blocks (**Figure 1;** Supplemental material **Table 1**). A 10 m transect tape was extended from each point consecutively in the south, west, north, east directions until twenty *M. capitata* colonies were found and tagged. Only corals that appeared healthy were tagged and sampled for this study.

**Figure 1.**
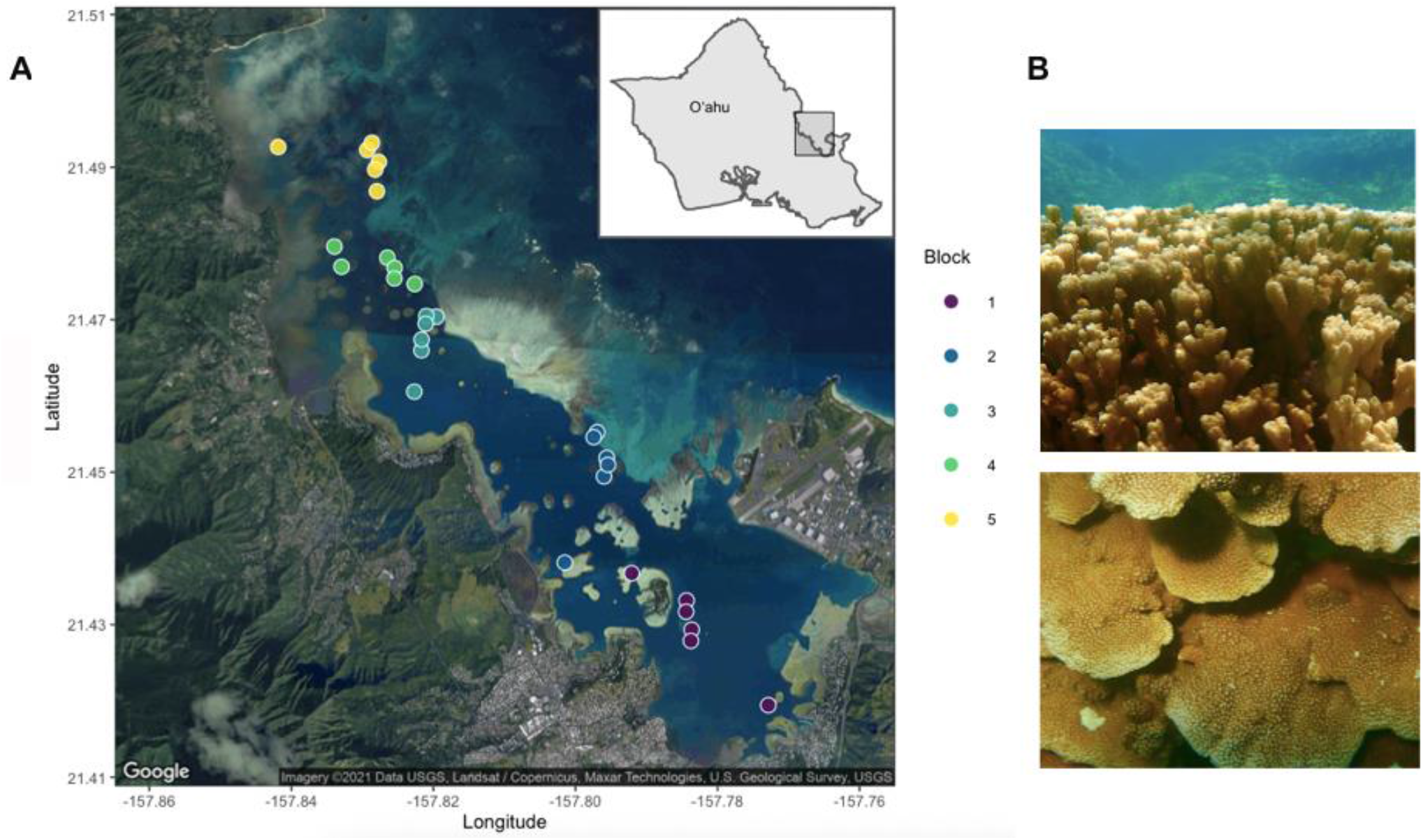
A. Sites and blocks in Kāneʻohe Bay. Each point is a randomly selected site within blocks represented by colors (Coordinates and site IDs in Supplemental material Table 1). Blocks go from the south bay (block 1) to north bay (block 5). B. *Montipora capitata*, the rice coral. The figures illustrate the morphological plasticity of this species, which can be branching (top) or plating (lower picture). Site IDs consist of the digit corresponding to the block in which the site is contained, followed by the site number (e.g., 1_10, with six sites per block, but not necessarily in consecutive order). Map done in the R package ggmap (Kahle & Wickham, 2013)

Temperature loggers (Hobo Pendant from Onset Computer Corp: UA-001-64 Data Logger) were wrapped in foil tape to prevent solar exposure (Bahr, Jokiel, & Rodgers, 2016) and attached to the cement block at the center of each site; the depth of the logger was assigned as the site depth. Temperature recordings every 10 min began on 12 July 2017 and continued until 26 July 2018, with the loggers periodically retrieved and recalibrated throughout the study period. Sediment traps were also attached to each cement block. These sediment traps were exchanged every 1-2 months, and the weight of sediment was used to estimate the sedimentation rate in each site following (Storlazzi, Field, & Bothner, 2011).

Early 2018, about 1cm^2^ was sampled from each of the 600 tagged *Montipora capitata* colonies. Sampled fragments were immediately preserved in 70% ethanol and stored at −20°C until processed. DNA was extracted using the Nucleospin Tissue Kits (Macherey-Nagel, Düren, Germany) following manufacturer instructions and quantified by fluorimetry (Quant-it HS dsDNA kit, Thermo-Fisher). Only corals that appeared healthy were sampled for this study.

### 2.2 SYMBIODINIACEAE ITS2 AMPLICON SEQUENCING LIBRARY PREPARATION

Symbiodiniaceae ITS2 amplicon library preparation and sequencing followed the protocol outlined in (Jacobs et al., 2021). Briefly, this involved amplification of the ITS2 region for each sample using individually barcoded algal symbiont-specific forward and reverse primers, that were pooled equimolarly, and prepared for sequencing using the KAPA hyper prep library kit (Roche/Kapa Biosystems, Cape Town, South Africa). Libraries were sequenced on the Illumina MiSeq platform (v3 2×300bp PE) at the University of Hawai‘i at Mānoa Advanced Studies in Genomic, Proteomic, and Bioinformatics sequencing facility.

### 2.3 AMPLICON SEQUENCING ANALYSIS WITH SYMPORTAL

Raw sequences were first demultiplexed using Cutadapt (Martin, 2011). Demultiplexed forward and reverse reads were submitted to SymPortal (Hume et al., 2019), a platform for genetically identifying Symbiodiniaceae using high through put ITS2 sequence data that differentiates intra- and intergenomic sources of *ITS2* sequence variance. After submission, sequence quality control is done within the SymPortal framework, using mothur 1.39.5 (Schloss et al., 2009), blast+ suite of executables (Camacho et al., 2009) and minimum entropy decomposition (MED; Eren et al. 2015). Sets of ITS2 sequences, occurring in a sufficient number of samples within both the dataset being analyzed and the entire database of samples run through SymPortal were identified as ‘defining intragenomic variants’ (DIVs) which were then used to characterize ITS2 type profiles.

Many terms have been used to describe the Symbiodiniaceae unity of resolution. To limit ambiguity, in this study, we will restrict our use to “Symbiodiniaceae type” and “Symbiodiniaceae profile”. A type refers to Symbiodiniaceae taxa that have a specific sequence as their most abundant sequence. A Symbiodiniaceae profile is a set of ITS2 sequences that have been found in a sufficient number of samples (DIV). For example, C17 is a Symbiodiniaceae type and C17d/C31-C21-C17e-C21ac-C17f-C17g is a Symbiodiniaceae profile with C17d present in higher abundance than the other types.

### 2.4 STATISTICAL ANALYSIS

All analyses were done in the R statistical environment (R Core Team 2020). Bray-Curtis dissimilarity of relative abundance of the Symbiodiniaceae community composition was tested by permutational multivariate analysis of variance (PERMANOVA) in the function *adonis* (for effect of block, with site nested within block), and pairwise.*adonis* (for pairwise PERMANOVA) in the vegan package (Dixon 2003), each with 999 permutations. To better visualize the similarity among blocks, R^2^ from the PERMANOVA was plotted in a dendrogram using the function *pheatmap* in the package gplots (Warnes et al. 2020). The function *metaMDS* was used in the R package *vegan* (Dixon 2003) to generate non-metric multidimensional scaling visualizations using Bray-Curtis dissimilarities of algal symbiont community per block.

Mean daily temperature, daily range and mean daily standard deviation were calculated per site using the R package *lubridate* (Grolemund & Wickham, 2011). The maximum site daily mean was considered as the maximum temperature, and the minimum site daily mean was considered the minimum temperature for the site.

Data loggers for 5 sites (2_2, 2_3, 3_2, 4_11 and 4_14) went missing during one deployment in the winter, so minimum and mean temperature for those sites were considered as the average minimum daily temperature of all sites, and the average daily temperature for all sites, respectively. The data logger for site 4_8 was lost during one deployment in summer, so maximum temperature and mean daily temperature for this site were replaced with the average maximum daily temperature of all sites, and the average daily temperature for all sites, respectively. Degree heating weeks (DHW) per site was calculated as the number of weeks when temperature exceeded the bleaching threshold of 28.5°C (MMM +1 °C; Wyatt et al. 2020; Dilworth et al. 2021.

To investigate the effect of the environmental data in driving the Symbiodiniaceae community, we performed distance-based redundancy analysis (dbRDA) using the function *capscale*. After running an ANOVA to check the significance of the constrains and the significance of the variables, we visualized the significant variables in the dbRDA. Samples were considered to have majority *Cladocopium* (C) or *Durusdinium* (D) if the proportion of either algal symbiont exceeded 80% in the sample (modified from Innis et al. 2018). All remaining samples with *Cladocopium* and *Durusdinium* were designated as CD, corresponding to corals with no dominant algal symbiont genus composing greater than 80% of reads. The relative contribution of each environmental factor in explaining the variation on the Symbiodiniaceae community was calculated as the total sum of squares for all variables divided by the sum of squares for that variable (Anderson 2017). Overall contribution of all environmental factors was calculated as the total sum of squares for all variables divided by the sum of residual and total sum of squares for all variables (Anderson 2017) Correlation among the environmental factors was calculated using the function *cor.test* in R. Data and R code to execute and reproduce all of the analyses and figures presented in the manuscript are archived at Zenodo (de Souza 2021).

## 3. Results

### 3.1 SYMBIODINIACEAE ITS2 SEQUENCES AND ITS2 TYPE PROFILES

From the 600 Montipora capitata sampled, 550 returned high quality Symbiodinaceae reads that passed initial quality control steps. Prior to quality filtering, the 550 samples returned 3,285,481 sequences and after sequence quality control and minimum entropy decomposition the SymPortal analysis returned 1,632,505 Symbiodiniaceae sequencing reads, an average of of 2,968 sequences per sample. Of those sequences, 11,386 were unique to our dataset and had not been previously identified by SymPortal. A total of 283 Symbiodiniacae types were identified, 85% belonging to genus *Cladocopium*, and 15% belonging to the genus *Durusdinium*. Twenty-six ITS2 type profiles were identified across all samples, twenty-three of which were from the genus *Cladocopium*, with the remaining 3 belonging to the genus *Durusdinium*. Overall, 43% of *Montipora capitata* hosted *Cladocopium* only, 11% hosted *Durusdinium* only, and 46% hosted a combination of both genera. From those mixed colonies, 32.5% were dominated by *Cladocopium* (C >80%), while 37% of colonies were dominated by *Durusdinium* (D>80%). Although it does not change the interpretation of results, to standardize the number of reads among samples, we excluded 20 samples whose number of reads were more than 2 standard deviations above or below the mean.

### 3.2 BIOGEOGRAPHY OF SYMBIODINIACEAE IN KĀNEʻOHE BAY

There is significant geographical structure in the Symbiodiniaceae composition of randomly sampled corals within each of the environmentally delineated blocks in Kāneʻohe Bay (PERMANOVA, F _25_= 6.5632, *p*= 0.001; **Table 1**). Pairwise comparisons revealed that blocks 1 and 5 were significantly different, whereas blocks in the middle of the bay (i.e., blocks 2-4) were rarely significantly different from one another (**Table 1**, **Figure 2A, B)**.

**Table 1.** PERMANOVA based on Bray-Curtis dissimilarities of the Symbiodiniaceae diversity present in corals sampled randomly from each environmentally defined block in Kāneʻohe Bay.

| Block | Df | Sum Sq | F | R<sup>2</sup> | P |
| --- | --- | --- | --- | --- | --- |
| Block | 4 | 2.8744 | 19.3604 | 0.10267 | 0.001* |
| Site: Block | 25 | 6.5632 | 7.0729 | 0.23443 | 0.001* |
| 1 vs 2 | 1 | 0.2667 | 4.9703 | 0.0243 | 0.17 |
| 1 vs 3 | 1 | 0.1488 | 2.9771 | 0.0148 | 0.65 |
| 1 vs 4 | 1 | 0.6422 | 12.5747 | 0.0575 | 0.01* |
| 1 vs 5 | 1 | 0.4994 | 13.3826 | 0.0612 | 0.01* |
| 2 vs 3 | 1 | 0.0328 | 0.5951 | 0.0028 | 1.00 |
| 2 vs 4 | 1 | 0.0900 | 1.6093 | 0.0073 | 1.00 |
| 2 vs 5 | 1 | 1.5508 | 36.1093 | 0.1432 | 0.01* |
| 3 vs 4 | 1 | 0.1997 | 3.7937 | 0.0173 | 0.45 |
| 3 vs 5 | 1 | 1.1910 | 30.1716 | 0.1235 | 0.01* |
| 4 vs 5 | 1 | 2.3907 | 58.4606 | 0.2077 | 0.01* |

**Figure 2.**
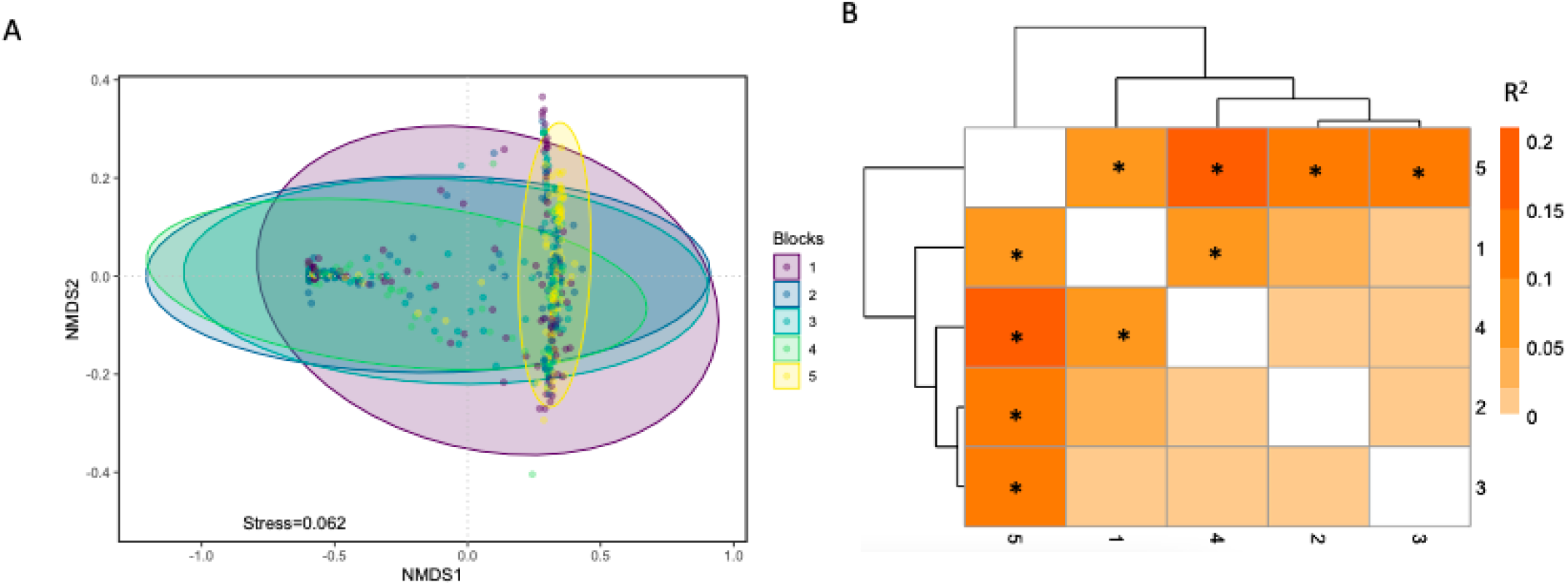
Similarity of Symbiodiniaceae communities detected. A) nMDS of Symbiodiniaceae per block. Ellipses are 95% confidence intervals. B) Dendrogram and heat map of R^2^ of the PERMANOVA of Symbiodiniaceae per block highlighting the similarity among blocks in the center of the bay.

Most sites had both *Cladocopium* and *Durusdinium* present, except two sites in block 5 (sites 5_3 and 5_6) which did not have any samples in which *Durusdinium* was detected (**Figure 3 a, b**; Supplemental material **Figure 1**). It is noteworthy that these two sites were the least diverse of all we sampled, with a few coral colonies hosting only one type of algal symbiont (C3), which was not seen in other sites (Supplemental material **Figure 1**). Corals within block 5 hosted significantly less *Durusdinium* symbionts than any other site, with all sites including corals hosting a majority of *Cladocopium*. Site 2_2 had the highest relative proportion of *Durusdinium* and site 3_2 was the only site to have three profiles of *Durusdinium*.

**Figure 3.**
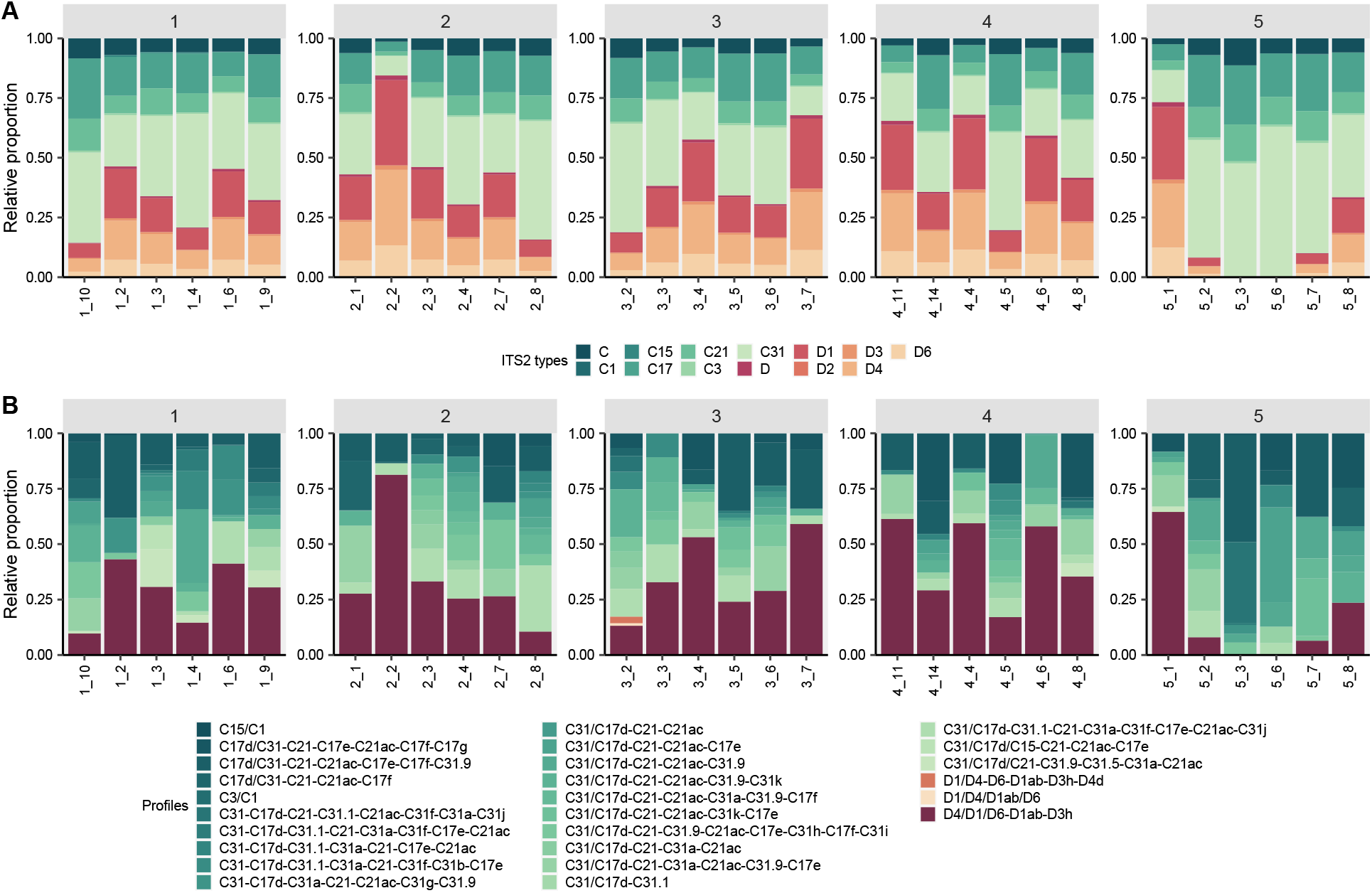
A. Major Symbiodiniaceae types by site and block. B) Symbiodiniaceae profiles by site. Block 1 is the southernmost in Kāneʻohe Bay, while block 5 is the most northern and similar to offshore conditions. Symbiodiniaceae ITS2 subtypes were summarized to the major subtype to facilitate visualization in the bar charts (i.e., C31a and C31b were summarized as C31). Due to the wide diversity of ITS2 available in the SymPortal database, not all sequences are given names. Only those sequences that are used in the definition of ITS2 type profiles (i.e. DIVs) are named. Unnamed *Cladocopium* and *Durusdinium* sequences were combined for visualization and represented as summed “C” and “D” types, respectively.

### 3.3 ENVIRONMENTAL DRIVERS OF SYMBIODINIACEAE COMMUNITY COMPOSITION

Depth of collection sites varied from 0.5 m to 3.5 m. Block 5 is the deepest block, with an average of 2.71 m, while sites in block 2 were shallowest, with an average of 1.36 m (Supplemental material **Table 1,2**). Sedimentation was highest at block 2 with an average of 0.67 g/day, and lowest at block 1 (0.03g/day). Average overall daily mean temperature in the bay for the year varied from 22.05 °C to 29.38°C. Block 1 and block 5 had lower daily temperature range and lower mean temperature standard deviation when compared to blocks in the center of the bay, which were more similar (Supplemental material **Table 2**; mean daily temperature range for block 1 and 5 were 0.24 °C and 0.74 °C respectively, while mean daily temperature range in the center of the bay varied from 1.24 °C to 1.87 °C; similarly, mean temperature standard deviation for block 1 and 5 were 0.24 °C and 0.20 °C respectively, while mean standard deviation in the center of the bay varied from 0.33 to 0.47).

dbRDA showed that *M. capitata* hosting majority *Cladocopium* are distinct from *M. capitata* hosting majority *Durusdinium* or mixed C & D (**Figure 4**). PERMANOVA of the environmental factors in the dbRDA **(Table 2),** showed that depth, DHW, maximum mean daily temperature recorded at the site, and mean daily standard deviation were all significant drivers of Symbiodiniaceae community composition. The environmental factors in the study explained only 20% of the Symbiodiniaceae variation, depth had the greatest relative contribution (61%) followed by daily standard deviation (19%), maximum temperature (9.9%) and DHW (4.8%) (**Table 2**).

**Figure 4.**
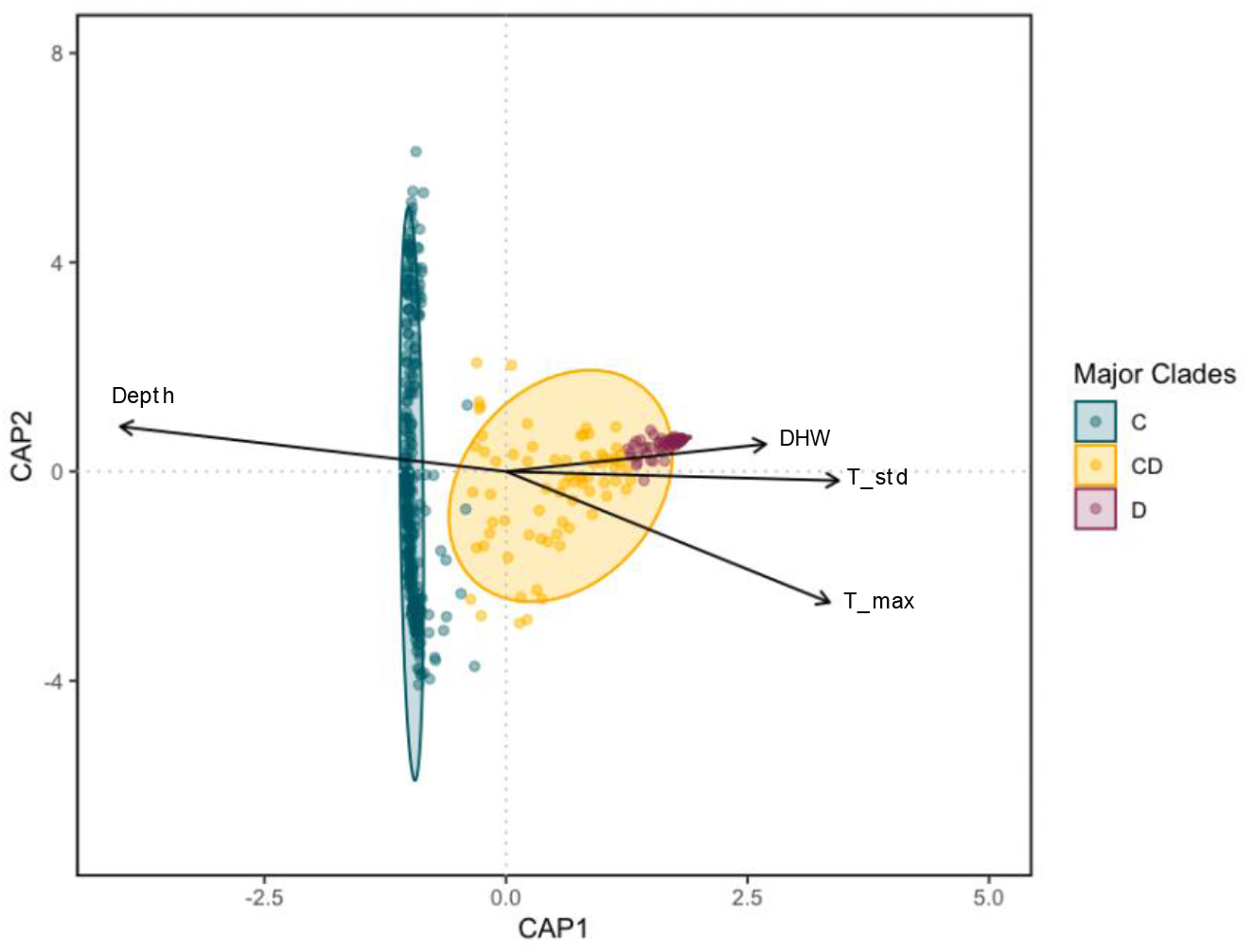
Distance based redundancy analysis (dbRDA) for environmental drivers of the Symbiodiniaceae communities measured in this study. Each point represents a *Montipora capitata* colony sampled irrespective of site. Samples were considered as majority *Cladocopium* (C) if they contain > 80%C, and majority *Durusdinium* (D) if > 80% D. Only vectors for the environmental factors contributing significantly to the algal symbiont diversity are plotted. Each arrow signifies the multiple partial correlation of the environmental driver in the RDA whose length and direction can be interpreted as indicative of its contribution to the explained variation.

**Table 2.** PERMANOVA of the environmental drivers of Bray-Curtis dissimilarities among Symbiodiniaceae communities among sites. Relative contribution was calculated as the sum square of each environmental factor divided by the sum of all environmental sum squares. Environmental factors explain 20% of Symbiodiniaceae variation.

| Environmental factors | Df | Sum Sq | F | P | Relative contribution |
| --- | --- | --- | --- | --- | --- |
| *Depth (m)* |  |  |  |  |  |
| Mean | 1 | 8.905 | 79.0332 | 0.001* | 0.610 |
| *Temperature (°C)* |  |  |  |  |  |
| DHW | 1 | 0.712 | 6.3152 | 0.009* | 0.0487 |
| Mean | 1 | 0.216 | 1.9169 | 0.149 | 0.0148 |
| Maximum | 1 | 1.448 | 12.8526 | 0.001* | 0.0992 |
| Minimum | 1 | 0.116 | 1.0312 | 0.314 | 0.0794 |
| Total range | 1 | 0.028 | 0.2523 | 0.810 | 0.0019 |
| Daily range | 1 | 0.029 | 0.2536 | 0.813 | 0.0019 |
| Mean daily standard deviation | 1 | 2.804 | 24.8804 | 0.001* | 0.1921 |
| *Sedimentation (g/day)* |  |  |  |  |  |
| Mean | 1 | 0.106 | 0.9397 | 0.339 | 0.0072 |
| Maximum | 1 | 0.025 | 0.2230 | 0.864 | 0.0017 |
| Minimum | 1 | 0.082 | 0.7298 | 0.435 | 0.0056 |
| Mean daily standard deviation | 1 | 0.123 | 1.0945 | 0.267 | 0.0084 |
| Residual |  | 58.255 |  |  |  |
| Total |  | 72.849 |  |  |  |

Most temperature parameters were significantly negatively correlated with depth (Supplemental material **Table 3**). Daily range and mean daily standard deviation were strongly inversely correlated to depth (r= −0.7425, p=<0.001; r= −0.7565, p= <0.001, respectively). In contrast, sedimentation parameters were weakly positively correlated with depth (r _548_=0.2380 to 0.2512, p=<0.001).

## 1. Discussion

Here, we contribute to a more nuanced understanding of the drivers of coral algal symbiont community composition by examining the associations of environmental drivers across a small spatial scale (~10km) on the community composition of Symbiodiniaceae in one of the dominant reef-building corals in the region, *Montipora capitata*.

Previous surveys of the Symbiodiniaceae community structure of *Montipora capitata* across Kāneʻohe Bay (Innis et al., 2018; Stat et al., 2011) found algal symbiont spatial structure at the level of site and colony, but not among different regions of the bay. Our finding of significant differences among blocks contrasts with the conclusions of Stat et al. (2011) that there was no regional structuring of algal symbiont communities. This difference may be a consequence of the way the bay was divided into regions in each study. In our study, sampling covered the entire bay, divided into 5 blocks based on the mean oceanographic conditions and water residence time, whereas Stat et al. (2011) restricted their comparisons to the south of the bay (which would equate to only blocks 1 and 2 in our study). However, we see significant differences even between blocks 1 and 2, so it may also be that our approach has greater statistical power. Stat et al. (2011) used cloning and Sanger sequencing of ITS2 to identify algal symbionts, so many fewer total colonies were sampled (52), each with far less resolution (5-7 clones per colony) than is possible with the amplicon approach using high throughput sequencing employed here.

Innis et al. (2018; Supplemental material **Figure 2**) sampled a large number of colonies (707) across the biophysical regions and depth range of Kāne‘ohe Bay, and like Stat et al. (2011), did not find evidence of Symbiodiniaceae regional differentiation across the environmental gradients of Kāneʻohe Bay. Innis et al. (2018) used a qPCR assay targeting *Cladocopium* and *Durusdinium* to show that orange morph colonies dominated by *Durusdinium* were more prevalent in shallow, high light environments (< 2 m), whereas brown-morph colonies, dominated by *Cladocopium*, were more prevalent with increasing depth and decreasing light. Although there was considerable variation, orange-morph colonies were more likely to be dominated by *Durusdinium* at shallow depths (< 4.3 m), and more likely dominated by *Cladocopium* below that depth, revealing a significant interaction between depth and Symbiodiniaceae community composition within the orange, but not the brown color morph. The authors concluded that Symbiodiniaceae community variability may arise from either holobiont phenotypic plasticity or differential survival of Symbiodiniaceae across light gradients and recommended additional study into whether algal symbiont communities were stable or plastic within individuals across these gradients (Innis et al. 2018). Although they collected random samples widely across the bay, they did not have the resolution to identify the diversity of subtypes of *Cladocopium* and *Durusdinium* as in this study and had no samples in the far north (block 5 in our study) and few at the far south (block 1) of the bay.

Using a metabarcoding approach, our study also shows that Symbiodiniaceae community composition varied at all spatial scales examined: among sites within a block, and among blocks across the bay. Most colonies (46%) hosted a mixture of both *Cladocopium* and *Durusdinium* variants, but nearly as many (43%) were dominated by only *Cladocopium* types, whereas only a few hosted exclusively *Durusdinium* types (11%). Interestingly, the relative proportion of algal symbiont types is relatively consistent among the colonies, no matter the location (i.e., when *Cladocopium* is present, C31 is typically most abundant followed by C17 and C21, whereas for *Durusdinium*, D1 is the most abundant, followed by D4 and D6), showing strong Symbiodiniaceae specificity in *Montipora capitata*. In total, seven types of *Cladocopium* (C1, C3, C15, C17, C21, C31) are detected among colonies across Kāneʻohe Bay, with C31, C17 and C1 being the most common *Cladocopium* types. Similarly, we detected six types of *Durusdinium* (D, D1, D2, D3, D4, D6) with D1, D4, and D6 being the most prevalent. These results are consistent with multiple studies reporting C31 to be the most common algae symbiont associated with *Montipora capitata* in Hawai‘i (Stat et al 2011; 2013; Cunning et al. 2016).

Despite considerable variation in the composition of the communities of Symbiodiniaceae hosted by *Montipora capitata* sampled across Kāneʻohe Bay, we find that the coral–algal symbiont association was strongly influenced by environmental gradients. *M. capitata* located at the environmental extremes (block 1 in the south and block 5 in the north) hosted Symbiodiniaceae communities that were significantly different from blocks in the center of the bay. We find 4 factors (depth, DHW, maximum and variability in temperature) that are significant drivers of Symbiodiniaceae community composition (**Table 2**), although, these 4 significant factors combined explained less than 20% of the variation in symbiont community composition. It is important to note that we cannot determine if these correlated environmental factors are the direct drivers of algal symbiont diversity. Until future studies determine the underlying mechanisms driving variation in algal symbiont community composition, we remain cautious in our interpretation of these environmental drivers.

It is interesting to note that corals that are dominated by *Durusdinium* are much closer together in the distance-based redundancy analysis (**Figure 4**), suggesting that corals hosting majority *Durusdinium* exist within a narrower set of environmental conditions, relative to a much wider suite of environmental conditions found for corals hosting majority *Cladocopium*. This is consistent with other studies that found *Durusdinium* to be present mostly in stressed colonies, because hosting *Durusdinium* has tradeoffs to the coral physiology and growth (Baker et al., 2013; Cunning et al., 2018; Matthews et al., 2018; Sproles et al., 2020, 2019)

### Depth

Consistent with previous studies (Baker, 2003; Innis et al., 2018; Rowan & Knowlton, 1995; Rowan et al., 1997; Toller, Rowan, & Knowlton, 2001), depth appears to be the strongest environmental driver of symbiont community composition measured in this study. The deepest block (5) hosted the lowest proportion of *Durusdinium*, with most colonies hosting only *Cladocopium* (**Figure 3**; Supplemental material **Figure 3)**. A similar pattern is found when looking at individual sites surveyed across all blocks, with the deepest sites (1_3, 1_4, 1_9, 1_10, 2_4 and 2_8) being mostly dominated by *Cladocopium*. While Innis et al. (2018) also found depth to be the primary driver of algal symbiont clades in Kāneʻohe Bay they concluded the mechanism was most likely light attenuation in deeper reefs as opposed to depth *per se*. It is well known that light attenuates with depth in water, but the relationship is complicated in coral reef environments where a variety of other factors can alter penetration of light to deeper corals (Edmunds, Tsounis, Boulon, & Bramanti, 2018; Hochberg, Peltier, & Maritorena, 2020; Storlazzi, Norris, & Rosenberger, 2015). Likewise, an entire suite of environmental parameters other than light also covary with depth, such that it is often difficult to know exactly which environmental or biological factors drive changes in community structure with depth (Kahng et al., 2010; Lesser, Slattery, & Leichter, 2009; Morgan, Moynihan, Sanwlani, & Switzer, 2020; Hoek, Breeman, Bak, & Buurt, 1978). Depth is simple to measure and is well-correlated with changes in coral reef community structure in many studies (reviewed by Kahng et al. 2010; Baldwin et al. 2018; Lesser et al. 2018) but is much harder to isolate as an environmental factor to determine the quantitative relative contribution in studies such as this.

### Sedimentation

Suspended sediments are one of those factors that can both impact corals directly and alter light penetration and irradiance of corals. Sedimentation can also impact corals negatively by increasing nutrient input, damaging coral surface and by making it harder for corals to feed and photosynthesize (Duckworth, Giofre, & Jones, 2017; Jones, Fisher, & Bessell-Browne, 2019; Weber et al., 2012).

Sediment particles in the water increase turbidity, which contributes to light attenuation, however, this might be affected by depth. Deeper lower sediment reefs (like most sites in block 5) may have higher light than some mid-range sites with higher turbidity. In our study, none of the sediment parameters were significant predictors of symbiont community composition (**Table 2**).

### Temperature

The role of temperature in mediating the symbiosis between the coral host and their algal endosymbionts has been widely studied and acknowledged as an important factor shaping Symbiodiniacae communities (Brown, 1997; Douglas, 2003; Fitt, Brown, Warner, & Dunne, 2001; Gates, Baghdasarian, & Muscatine, 1992), so it is not surprising that several aspects of temperature come out as significant drivers of community composition. Degree heating weeks (DHW) and maximum temperature are widely established to be major predictors of the breakdown of symbiosis between the partners and result in coral bleaching (Sully et al., 2019; Thompson & Woesik, 2009). Unsurprisingly, both were significant in our analysis of environmental drivers, although neither explained much of the variation in algal symbiont community structure among sites. In contrast, variability in temperature, in this case mean daily temperature standard deviation, had the second largest contribution to the observed variation in algal symbiont community structure. A number of studies highlight the importance of daily temperature fluctuations for coral acclimatization to higher temperatures (Barshis et al., 2010; Safaie et al., 2018), suggesting that temperature fluctuations encourage greater thermal tolerance by exposing the corals to short periods of thermal stress without causing mortality. Blocks in the center of the bay had higher temperature variation (higher mean average temperature standard deviation and higher mean daily range) and our data shows that the existing community structure of algal symbionts responds to such variability (**Figure 4**). Blocks in the center of the bay (blocks 2-4) had higher proportion of *Durusdinium*, while block 1 and block 5 (extreme north and south, respectively) had the lowest proportion, suggesting that blocks at the extreme ends of the bay may be more vulnerable to bleaching events. This is consistent with the findings of (Bahr, Rodgers, & Jokiel, 2017), which surveyed corals during the 2015 bleaching event in Kāneʻohe Bay and found that the highest levels of bleaching and paling happened in the north of the bay (70%) with 18% mortality, whereas the highest death rate was observed in the south of the bay (28%) despite somewhat lower (60%) bleaching and paling.

While hosting *Durusdinium* can increase the survival of corals under elevated temperatures (Berkelmans & van Oppen, 2006), thermal tolerance often comes at a cost, because it also tends to decrease growth, reproduction, and metabolite rate exchange with the host (Baker et al., 2013; Cunning et al., 2018; Jones & Berkelmans, 2010; Jones & Berkelmans, 2011; Matthews et al., 2018; Sproles et al., 2020, 2019). While there are patterns at the level of genera, recent studies also highlight the need to look at finer scale identification, because examination of subtypes (some of which remain undescribed and some of which are now named species, LaJeunesse et al. 2018) has unraveled biologically relevant patterns such as host specificity, niche diversification, and physiological differences between algal symbiont subtypes (Iglesias-Prieto, Beltrán, LaJeunesse, Reyes-Bonilla, & Thomé, 2004; Sampayo, Ridgway, Bongaerts, & Hoegh-Guldberg, 2008).

Here we advance the previous work of Stat et al. (2011) and Innis et al. (2018) by showing that algal symbiont communities within a single species of coral in a single embayment are finely tuned to their environmental conditions. Whether this results from selection for Symbiodiniaceae types living under different environments, or adaptive shuffling of Symbiodiniaceae communities in response to environmental conditions or both, remains to be determined. Further, as outlined above, both depth and sedimentation may have complex interactions with light that appear to play a role in determining algal symbiont community composition as well. However, an important caveat of this study is that light was not measured, and each site is characterized by a central point, rather than fine-scale measurements of environmental drivers at the level of individual colonies. A more detailed understanding of the relationship between adaptive tuning of algal symbiont communities to the environmental conditions under which the holobiont is living will require fine-scale environmental measurements coupled with long-term monitoring of corals in the field to determine whether and how algal symbiont communities within individual colonies change through time. Comparing microscale environmental variability (Gorospe & Karl, 2011, 2015) to algal symbiont community structure might explain much of the variability we see, because algal symbionts in colonies may be adaptively responding to fine-scale variability at the same site within the broad regional differences we compared here.

### Conclusions

Fine-scale sampling of 600 *M. capitata* colonies across a relatively small spatial gradient (~10km) within Kāneʻohe Bay showed that biogeographical patterns of algal symbiont distribution are driven by environmental variables associated with depth, temperature, and sedimentation. This fine-scale variation in algal symbiont communities across environmental gradients suggests that algal symbiont communities appear to adaptively match the environmental conditions surrounding the holobiont.

Our study highlights the complex interactions among environmental factors and algal symbiont diversity in the reef-building coral *Montipora capitata*. While depth was the main factor driving algal symbiont community composition in our study, other factors also contributed to the algal symbiont diversity, including minimum sedimentation and various measurements of temperature (DHW, max temp & mean standard deviation). We also note that many factors correlate with depth, such as light, temperature, sedimentation rate, and water flow, such that fine-scale measurements of the full range of environmental factors surrounding individual colonies through time will be needed to pinpoint the most important environmental drivers of algal symbiont community structure. Regardless of the ultimate suite of parameters that drive algal symbiont community structure in corals, our study shows that Symbiodiniaceae communities are attuned to fine-scale environmental gradients and that understanding these complex interactions across the heterogeneous mosaic of coral reef environments is needed to better predict spatial patterns in biological responses such as bleaching susceptibility.

## Acknowledgments

All members of the Coral Resilience Lab (the legacy of Ruth Gates) and ToBo Lab who assisted with collections, molecular work, or bioinformatics (special thanks to Kira Hughes, Joshua Hancock, Janaya Bruce, Dennis Conneta, Caroline Hobbs, Valerie Kahkejian, and Christian Marin). This work was funded by a Coordenação de Aperfeiçoamento de Pessoal de Nível Superior – Brasil (CAPES) fellowship, the National Science Foundation (OA-1416889 & BioOCE-1924604 to RJT) and the Paul G. Allen Family Foundation. This is SOEST contribution number xxx and HIMB contribution number xxx.

## Compliance with ethical standards

Conflict of interest: On behalf of all authors, the corresponding author states that there is no conflict of interest.

## Contributions

M.R.S, C.C., L.R.J., R.G. conceived of and designed the experiment, and M.R.S, C.C. & L.R.J. led the collections. M.R.S, C.C., L.R.J. contributed to molecular work. M.R.S., C.D., R.J.T. contributed to analysis; C.D., R.G., R.J.T. contributed to funding. The first draft of the paper was written by M.R.S with help from R.J.T. All authors contributed to discussion, data interpretation, manuscript revisions, and all approve the final manuscript.

## References

Anderson MJ (2017) Permutational multivariate analysis of variance (PERMANOVA). Wiley StatsRef: Statistics Reference Online. Wiley Online Library, eds Balakrishnan N, et al. Available at https://onlinelibrary.wiley.com/doi/10.1002/9781118445112.stat07841

Bahr, K. D., Jokiel, P. L., & Rodgers, K. S. (2016). Influence of solar irradiance on underwater temperature recorded by temperature loggers on coral reefs: Evaluation of underwater temperature loggers. Limnology and Oceanography: Methods, 14(5), 338–342. doi: 10.1002/lom3.10093

Bahr, K. D., Jokiel, P. L., & Toonen, R. J. (2015). The unnatural history of Kāne‘ohe Bay: Coral reef resilience in the face of centuries of anthropogenic impacts. PeerJ, 3, e950. doi: 10.7717/peerj.950

Bahr, K. D., Rodgers, K. S., & Jokiel, P. L. (2017). impact of three bleaching events on the reef resiliency of Kāne‘ohe Bay, Hawai‘i. Frontiers in Marine Science, 4, 398. doi: 10.3389/fmars.2017.00398

Baird, A., Cumbo, V., Leggat, W., & Rodriguez-Lanetty, M. (2007). Fidelity and flexibility in coral symbioses. Marine Ecology Progress Series, 347, 307–309. doi: 10.3354/meps07220

Baker, A. C. (2003). Flexibility and specificity in coral-algal symbiosis: diversity, ecology, and biogeography of *Symbiodinium*. Annual Review of Ecology, Evolution, and Systematics, 34(1), 661–689. doi: 10.1146/annurev.ecolsys.34.011802.132417

Baker, D. M., Andras, J. P., Jordán-Garza, A. G., & Fogel, M. L. (2013). Nitrate competition in a coral symbiosis varies with temperature among *Symbiodinium* clades. The ISME Journal, 7(6), 1248–1251. doi: 10.1038/ismej.2013.12

Baldwin, C. C., Tornabene, L., & Robertson, D. R. (2018). Below the mesophotic. Scientific Reports, 8(1), 4920. doi: 10.1038/s41598-018-23067-1

Barshis, D. J., Stillman, J. H., Gates, R. D., Toonen, R. J., Smith, L. W., & Birkeland, C. (2010). Protein expression and genetic structure of the coral *Porites lobata* in an environmentally extreme Samoan back reef: Does host genotype limit phenotypic plasticity? Molecular Ecology, 19(8), 1705–1720. doi: 10.1111/j.1365-294X.2010.04574.x

Berkelmans, R., & van Oppen, M. J. H. (2006). The role of zooxanthellae in the thermal tolerance of corals: A ‘nugget of hope’ for coral reefs in an era of climate change. Proceedings of the Royal Society B: Biological Sciences, 273(1599), 2305–2312. doi: 10.1098/rspb.2006.3567

Bongaerts, P., Ridgway, T., Sampayo, E. M., & Hoegh-Guldberg, O. (2010). Assessing the ‘deep reef refugia’ hypothesis: Focus on Caribbean reefs. Coral Reefs, 29(2), 309–327. doi: 10.1007/s00338-009-0581-x

Brown, B. E. (1997). Coral bleaching: Causes and consequences. Coral Reefs, 16(0), S129–S138. doi: 10.1007/s003380050249

Buddemeier, R. W., & Fautin, D. G. (1993). Coral bleaching as an adaptive mechanism. BioScience, 43(5), 320–326. doi: 10.2307/1312064

Camacho, C., Coulouris, G., Avagyan, V., Ma, N., Papadopoulos, J., Bealer, K., & Madden, T. L. (2009). BLAST+: Architecture and applications. BMC Bioinformatics, 10(1), 421. doi: 10.1186/1471-2105-10-421

Caruso, C., de Souza, M. R., Ruiz-Jones, L., Conetta, D., Hancock, J., Hobbs, C., … Drury, C. (2021). Genetic patterns in *Montipora capitata* across an environmental mosaic in Kāne’ohe Bay. doi: 10.1101/2021.10.07.463582

Cunning, R, Ritson-Williams, R., & Gates, R. (2016). Patterns of bleaching and recovery of Montipora capitata in Kāne‘ohe Bay, Hawai‘i, USA. Marine Ecology Progress Series, 551, 131–139. doi: 10.3354/meps11733

Cunning, Ross, Silverstein, R. N., & Baker, A. C. (2018). Symbiont shuffling linked to differential photochemical dynamics of *Symbiodinium* in three Caribbean reef corals. Coral Reefs, 37(1), 145–152. doi: 10.1007/s00338-017-1640-3

Dilworth, J., Caruso, C., Kahkejian, V. A., Baker, A. C., & Drury, C. (2021). Host genotype and stable differences in algal symbiont communities explain patterns of thermal stress response of Montipora capitata following thermal pre-exposure and across multiple bleaching events. Coral Reefs, 40(1), 151–163. doi: 10.1007/s00338-020-02024-3

de Souza, M.R. (2021). Community composition of coral-associated Symbiodiniaceae is driven by fine-scale environmental gradients. https://doi.org/10.5281/zenodo.5670832

Dixon, P. (2003). VEGAN, a package of R functions for community ecology. Journal of Vegetation Science: Official Organ of the International Association for Vegetation Science, 14(6), 927–930. doi: 10.1111/j.1654-1103.2003.tb02228.x

Donner, S. D., Skirving, W. J., Little, C. M., Oppenheimer, M., & Hoegh-Guldberg, O. (2005). Global assessment of coral bleaching and required rates of adaptation under climate change. Global Change Biology, 11(12), 2251–2265. doi: 10.1111/j.1365-2486.2005.01073.x

Douglas, A. E. (2003). Coral bleaching––how and why? Marine Pollution Bulletin, 46(4), 385–392. doi: 10.1016/S0025-326X(03)00037-7

Duckworth, A., Giofre, N., & Jones, R. (2017). Coral morphology and sedimentation. Marine Pollution Bulletin, 125(1–2), 289–300. doi: 10.1016/j.marpolbul.2017.08.036

Edmunds, P. J., Tsounis, G., Boulon, R., & Bramanti, L. (2018). Long-term variation in light intensity on a coral reef. Coral Reefs, 37(3), 955–965. doi: 10.1007/s00338-018-1721-y

Eren, A. M., Morrison, H. G., Lescault, P. J., Reveillaud, J., Vineis, J. H., & Sogin, M. L. (2015). Minimum entropy decomposition: Unsupervised oligotyping for sensitive partitioning of high-throughput marker gene sequences. The ISME Journal, 9(4), 968–979. doi: 10.1038/ismej.2014.195

Fautin, D. G., & Buddemeier, R. W. (2004). Adaptive bleaching: A general phenomenon. Hydrobiologia, 530/531, 459–467.

Finney, J. C., Pettay, D. T., Sampayo, E. M., Warner, M. E., Oxenford, H. A., & LaJeunesse, T. C. (2010). The relative significance of host–habitat, depth, and geography on the ecology, endemism, and speciation of coral endosymbionts in the genus *Symbiodinium*. Microbial Ecology, 60(1), 250–263. doi: 10.1007/s00248-010-9681-y

Fitt, W., Brown, B., Warner, M., & Dunne, R. (2001). Coral bleaching: Interpretation of thermal tolerance limits and thermal thresholds in tropical corals. Coral Reefs, 20(1), 51–65. doi: 10.1007/s003380100146

Gates, R. D., Baghdasarian, G., & Muscatine, L. (1992). Temperature stress causes host cell detachment in symbiotic cnidarians: implications for coral bleaching. The Biological Bulletin, 182(3), 324–332. doi: 10.2307/1542252

Gorospe, K. D., & Karl, S. A. (2011). Small-scale spatial analysis of *in situ* sea temperature throughout a single coral patch reef. Journal of Marine Biology, 2011, 1–12. doi: 10.1155/2011/719580

Gorospe, K. D., & Karl, S. A. (2015). Depth as an organizing force in *Pocillopora damicornis*: intra-reef genetic architecture. PLOS ONE, 10(3), e0122127. doi: 10.1371/journal.pone.0122127

Goulet, T. (2006). Most corals may not change their symbionts. Marine Ecology Progress Series, 321, 1–7. doi: 10.3354/meps321001

Goulet, T. (2007). Most scleractinian corals and octocorals host a single symbiotic zooxanthella clade. Marine Ecology Progress Series, 335, 243–248. doi: 10.3354/meps335243

Goulet, T., & Coffroth, M. (2003). Stability of an octocoral-algal symbiosis over time and space. Marine Ecology Progress Series, 250, 117–124. doi: 10.3354/meps250117

Grégoire, V., Schmacka, F., Coffroth, M. A., & Karsten, U. (2017). Photophysiological and thermal tolerance of various genotypes of the coral endosymbiont *Symbiodinium* sp. (Dinophyceae). Journal of Applied Phycology, 29(4), 1893–1905. doi: 10.1007/s10811-017-1127-1

Grolemund, G., & Wickham, H. (2011). Dates and times made easy with lubridate. Journal of Statistical Software, 40(3). doi: 10.18637/jss.v040.i03

Heron, S. F., Maynard, J. A., van Hooidonk, R., & Eakin, C. M. (2016). Warming trends and bleaching stress of the world’s coral reefs 1985–2012. Scientific Reports, 6(1), 38402. doi: 10.1038/srep38402

Hicks, C. C., Graham, N. A. J., & Cinner, J. E. (2013). Synergies and tradeoffs in how managers, scientists, and fishers value coral reef ecosystem services. Global Environmental Change, 23(6), 1444–1453. doi: 10.1016/j.gloenvcha.2013.07.028

Hoadley, K. D., Pettay, D. T., Grottoli, A. G., Cai, W.-J., Melman, T. F., Schoepf, V., … Warner, M. E. (2015). Physiological response to elevated temperature and pCO2 varies across four Pacific coral species: Understanding the unique host+symbiont response. Scientific Reports, 5(1), 18371. doi: 10.1038/srep18371

Hochberg, E. J., Peltier, S. A., & Maritorena, S. (2020). Trends and variability in spectral diffuse attenuation of coral reef waters. Coral Reefs, 39(5), 1377–1389. doi: 10.1007/s00338-020-01971-1

Hoegh-Guldberg, O., Mumby, P. J., Hooten, A. J., Steneck, R. S., Greenfield, P., Gomez, E., … Hatziolos, M. E. (2007). Coral reefs under rapid climate change and ocean acidification. Science, 318(5857), 1737–1742. doi: 10.1126/science.1152509

Hoegh-Guldberg, Ove. (1999). Climate change, coral bleaching and the future of the world’s coral reefs. Marine and Freshwater Research. doi: 10.1071/MF99078

Hoegh-Guldberg, Ove, Jones, R. J., Ward, S., & Loh, W. K. (2002). Is coral bleaching really adaptive? Nature, 415(6872), 601–602. doi: 10.1038/415601a

Howells, E. J., Beltran, V. H., Larsen, N. W., Bay, L. K., Willis, B. L., & van Oppen, M. J. H. (2012). Coral thermal tolerance shaped by local adaptation of photosymbionts. Nature Climate Change, 2(2), 116–120. doi: 10.1038/nclimate1330

Hughes, T. P. (2003). Climate change, human impacts, and the resilience of coral reefs. Science, 301(5635), 929–933. doi: 10.1126/science.1085046

Hughes, Terry P., Kerry, J. T., Álvarez-Noriega, M., Álvarez-Romero, J. G., Anderson, K. D., Baird, A. H., … Wilson, S. K. (2017). Global warming and recurrent mass bleaching of corals. Nature, 543(7645), 373–377. doi: 10.1038/nature21707

Hume, B. C. C., Smith, E. G., Ziegler, M., Warrington, H. J. M., Burt, J. A., LaJeunesse, T. C., … Voolstra, C. R. (2019). SymPortal: A novel analytical framework and platform for coral algal symbiont next-generation sequencing *ITS2* profiling. Molecular Ecology Resources, 19(4), 1063–1080. doi: 10.1111/1755-0998.13004

Iglesias-Prieto, R., Beltrán, V. H., LaJeunesse, T. C., Reyes-Bonilla, H., & Thomé, P. E. (2004). Different algal symbionts explain the vertical distribution of dominant reef corals in the eastern Pacific. Proceedings of the Royal Society of London. Series B: Biological Sciences, 271(1549), 1757–1763. doi: 10.1098/rspb.2004.2757

Innis, T., Cunning, R., Ritson-Williams, R., Wall, C. B., & Gates, R. D. (2018). Coral color and depth drive symbiosis ecology of Montipora capitata in Kāne‘ohe Bay, O‘ahu, Hawai‘i. Coral Reefs, 37(2), 423–430. doi: 10.1007/s00338-018-1667-0

Jacobs, K. P., Hunter, C. L., Forsman, Z. H., Pollock, A. L., de Souza, M. R., & Toonen, R. J. (2021). A phylogenomic examination of Palmyra Atoll’s corallimorpharian invader. Coral Reefs. doi: 10.1007/s00338-021-02143-5

Jones, A., & Berkelmans, R. (2010). Potential costs of acclimatization to a warmer climate: growth of a reef coral with heat tolerant vs. sensitive symbiont types. PLoS ONE, 5(5), e10437. doi: 10.1371/journal.pone.0010437

Jones, A. M., & Berkelmans, R. (2011). Tradeoffs to thermal acclimation: energetics and reproduction of a reef coral with heat tolerant *Symbiodinium* type-D. Journal of Marine Biology, 2011, 1–12. doi: 10.1155/2011/185890

Jones, R., Fisher, R., & Bessell-Browne, P. (2019). Sediment deposition and coral smothering. PLOS ONE, 14(6), e0216248. doi: 10.1371/journal.pone.0216248

Kahle, D., & Wickham, H. (2013). ggmap: Spatial Visualization with ggplot2. The R Journal, 5(1), 144. doi: 10.32614/RJ-2013-014

Kahng, S. E., Garcia-Sais, J. R., Spalding, H. L., Brokovich, E., Wagner, D., Weil, E., … Toonen, R. J. (2010). Community ecology of mesophotic coral reef ecosystems. Coral Reefs, 29(2), 255–275. doi: 10.1007/s00338-010-0593-6

LaJeunesse, T. C. (2005). “Species” radiations of symbiotic dinoflagellates in the Atlantic and Indo-Pacific since the Miocene-Pliocene transition. Molecular Biology and Evolution, 22(3), 570–581. doi: 10.1093/molbev/msi042

LaJeunesse, T. C., Parkinson, J. E., Gabrielson, P. W., Jeong, H. J., Reimer, J. D., Voolstra, C. R., & Santos, S. R. (2018). Systematic revision of Symbiodiniaceae highlights the antiquity and diversity of coral endosymbionts. Current Biology, 28(16), 2570–2580.e6. doi: 10.1016/j.cub.2018.07.008

LaJeunesse, T. C., Pettay, D. T., Sampayo, E. M., Phongsuwan, N., Brown, B., Obura, D. O., … Fitt, W. K. (2010). Long-standing environmental conditions, geographic isolation and host-symbiont specificity influence the relative ecological dominance and genetic diversification of coral endosymbionts in the genus *Symbiodinium*. Journal of Biogeography, 37(5), 785–800. doi: 10.1111/j.1365-2699.2010.02273.x

LaJeunesse, ToddC., Thornhill, DanielJ., Cox, EvelynF., Stanton, FrankG., Fitt, WilliamK., & Schmidt, GregoryW. (2004). High diversity and host specificity observed among symbiotic dinoflagellates in reef coral communities from Hawaii. Coral Reefs. doi: 10.1007/s00338-004-0428-4

Lesser, M. P., Slattery, M., & Leichter, J. J. (2009). Ecology of mesophotic coral reefs. Journal of Experimental Marine Biology and Ecology, 375(1–2), 1–8. doi: 10.1016/j.jembe.2009.05.009

Lesser, M. P., Slattery, M., & Mobley, C. D. (2018). Biodiversity and functional ecology of mesophotic coral reefs. Annual Review of Ecology, Evolution, and Systematics, 49(1), 49–71. doi: 10.1146/annurev-ecolsys-110617-062423

Lough, J. M., Anderson, K. D., & Hughes, T. P. (2018). Increasing thermal stress for tropical coral reefs: 1871–2017. Scientific Reports, 8(1), 6079. doi: 10.1038/s41598-018-24530-9

Lowe, R. J., Falter, J. L., Monismith, S. G., & Atkinson, M. J. (2009a). A numerical study of circulation in a coastal reef-lagoon system. Journal of Geophysical Research, 114(C6), C06022. doi: 10.1029/2008JC005081

Lowe, R. J., Falter, J. L., Monismith, S. G., & Atkinson, M. J. (2009b). Wave-driven circulation of a coastal reef–lagoon system. Journal of Physical Oceanography, 39(4), 873–893. doi: 10.1175/2008JPO3958.1

Martin, M. (2011). Cutadapt removes adapter sequences from high-throughput sequencing reads. EMBnet.Journal, 17(1), 3. doi: https://doi.org/10.14806/ej.17.1.200

Masson-Delmotte, V., Zhai, P., Pirani, A., Connors, S. L., Péan, C., Berger, S., …Zhou, B. (2021) IPCC, 2021: Climate Change 2021: The Physical science basis. Contribution of working group I to the sixth assessment report of the intergovernmental panel on climate change (eds.)]. Cambridge University Press. In Press.

Matthews, J. L., Oakley, C. A., Lutz, A., Hillyer, K. E., Roessner, U., Grossman, A. R., … Davy, S. K. (2018). Partner switching and metabolic flux in a model cnidarian–dinoflagellate symbiosis. Proceedings of the Royal Society B: Biological Sciences, 285(1892), 20182336. doi: 10.1098/rspb.2018.2336

McIlroy, S. E., Gillette, P., Cunning, R., Klueter, A., Capo, T., Baker, A. C., & Coffroth, M. A. (2016). The effects of *Symbiodinium* (Pyrrhophyta) identity on growth, survivorship, and thermal tolerance of newly settled coral recruits. Journal of Phycology, 52(6), 1114–1124. doi: 10.1111/jpy.12471

Moberg, F., & Folke, C. (1999). Ecological goods and services of coral reef ecosystems. Ecological Economics, 29(2), 215–233. doi: 10.1016/S0921-8009(99)00009-9

Morgan, K. M., Moynihan, M. A., Sanwlani, N., & Switzer, A. D. (2020). Light limitation and depth-variable sedimentation drives vertical reef compression on turbid coral reefs. Frontiers in Marine Science, 7, 571256. doi: 10.3389/fmars.2020.571256

Muscatine, L., & Porter, J. W. (1977). Reef corals: mutualistic symbioses adapted to nutrient-poor environments. BioScience, 27(7), 454–460. doi: 10.2307/1297526

Oliver, J. K., Berkelmans, R., & Eakin, C. M. (2018). Coral bleaching in space and time. In M. J. H. van Oppen & J. M. Lough (Eds.), Coral bleaching: patterns, processes, causes and consequences (pp. 27–49). Cham: Springer International Publishing. doi: 10.1007/978-3-319-75393-5_3

Oliver, T. A., & Palumbi, S. R. (2011). Do fluctuating temperature environments elevate coral thermal tolerance? Coral Reefs, 30(2), 429–440. doi: 10.1007/s00338-011-0721-y

Peñaflor, E. L., Skirving, W. J., Strong, A. E., Heron, S. F., & David, L. T. (2009). Sea-surface temperature and thermal stress in the Coral Triangle over the past two decades. Coral Reefs, 28(4), 841–850. doi: 10.1007/s00338-009-0522-8

Quigley, K. M., Baker, A. C., Coffroth, M. A., Willis, B. L., & van Oppen, M. J. H. (2018). Bleaching resistance and the role of algal endosymbionts. In M. J. H. van Oppen & J. M. Lough (Eds.), Coral Bleaching (pp. 111–151). Cham: Springer International Publishing. doi: 10.1007/978-3-319-75393-5_6

Robison, J. D., & Warner, M. E. (2006). Differential impacts of photoacclimation and thermal stress on the photobiology of four different phylotypes of *Symbiodinium* (Pyrrhophyta). Journal of Phycology, 42(3), 568–579. doi: 10.1111/j.1529-8817.2006.00232.x

Rowan, R., & Knowlton, N. (1995). Intraspecific diversity and ecological zonation in coral-algal symbiosis. Proceedings of the National Academy of Sciences, 92(7), 2850–2853. doi: 10.1073/pnas.92.7.2850

Rowan, Rob. (2004). Thermal adaptation in reef coral symbionts. Nature, 430(7001), 742–742. doi: 10.1038/430742a

Rowan, Rob, Knowlton, N., Baker, A., & Jara, J. (1997). Landscape ecology of algal symbionts creates variation in episodes of coral bleaching. Nature, 388(6639), 265–269. doi: 10.1038/40843

Safaie, A., Silbiger, N. J., McClanahan, T. R., Pawlak, G., Barshis, D. J., Hench, J. L., … Davis, K. A. (2018). High frequency temperature variability reduces the risk of coral bleaching. Nature Communications, 9(1), 1671. doi: 10.1038/s41467-018-04074-2

Sampayo, E. M., Ridgway, T., Bongaerts, P., & Hoegh-Guldberg, O. (2008). Bleaching susceptibility and mortality of corals are determined by fine-scale differences in symbiont type. Proceedings of the National Academy of Sciences, 105(30), 10444–10449. doi: 10.1073/pnas.0708049105

Schloss, P. D., Westcott, S. L., Ryabin, T., Hall, J. R., Hartmann, M., Hollister, E. B., … Weber, C. F. (2009). Introducing mothur: Open-Source, Platform-Independent, Community-Supported Software for Describing and Comparing Microbial Communities. Applied and Environmental Microbiology, 75(23), 7537–7541. doi: 10.1128/AEM.01541-09

Sproles, A. E., Oakley, C. A., Krueger, T., Grossman, A. R., Weis, V. M., Meibom, A., & Davy, S. K. (2020). Sub-cellular imaging shows reduced photosynthetic carbon and increased nitrogen assimilation by the non-native endosymbiont *Durusdinium trenchii* in the model cnidarian Aiptasia. Environmental Microbiology, 22(9), 3741–3753. doi: 10.1111/1462-2920.15142

Sproles, A. E., Oakley, C. A., Matthews, J. L., Peng, L., Owen, J. G., Grossman, A. R., … Davy, S. K. (2019). Proteomics quantifies protein expression changes in a model cnidarian colonised by a thermally tolerant but suboptimal symbiont. The ISME Journal, 13(9), 2334–2345. doi: 10.1038/s41396-019-0437-5

Stat, M., Bird, C. E., Pochon, X., Chasqui, L., Chauka, L. J., Concepcion, G. T., … Gates, R. D. (2011). Variation in *Symbiodinium* ITS2 sequence assemblages among coral colonies. PLoS ONE, 6(1), e15854. doi: 10.1371/journal.pone.0015854

Stat, M., & Gates, R. D. (2011). Clade D *Symbiodinium* in Scleractinian corals: a “nugget” of hope, a selfish opportunist, an ominous sign, or all of the above? Journal of Marine Biology, 2011, 1–9. doi: 10.1155/2011/730715

Stat, M., Pochon, X., Franklin, E. C., Bruno, J. F., Casey, K. S., Selig, E. R., & Gates, R. D. (2013). The distribution of the thermally tolerant symbiont lineage (*Symbiodinium* clade D) in corals from Hawaii: Correlations with host and the history of ocean thermal stress. Ecology and Evolution, 3(5), 1317–1329. doi: 10.1002/ece3.556

Storlazzi, C. D., Field, M. E., & Bothner, M. H. (2011). The use (and misuse) of sediment traps in coral reef environments: Theory, observations, and suggested protocols. Coral Reefs, 30(1), 23–38. doi: 10.1007/s00338-010-0705-3

Storlazzi, Curt D., Norris, B. K., & Rosenberger, K. J. (2015). The influence of grain size, grain color, and suspended-sediment concentration on light attenuation: Why fine-grained terrestrial sediment is bad for coral reef ecosystems. Coral Reefs, 34(3), 967–975. doi: 10.1007/s00338-015-1268-0

Sully, S., Burkepile, D. E., Donovan, M. K., Hodgson, G., & van Woesik, R. (2019). A global analysis of coral bleaching over the past two decades. Nature Communications, 10(1), 1264. doi: 10.1038/s41467-019-09238-2

Thompson, D. M., & van Woesik, R. (2009). Corals escape bleaching in regions that recently and historically experienced frequent thermal stress. Proceedings of the Royal Society B: Biological Sciences, 276(1669), 2893–2901. doi: 10.1098/rspb.2009.0591

Toller, W. W., Rowan, R., & Knowlton, N. (2001). Repopulation of zooxanthellae in the caribbean corals *Montastraea annularis* and *M. faveolata* following experimental and disease-associated bleaching. The Biological Bulletin, 201(3), 360–373. doi: 10.2307/1543614

Tonk, L., Sampayo, E. M., Weeks, S., Magno-Canto, M., & Hoegh-Guldberg, O. (2013). Host-specific interactions with environmental factors shape the distribution of *Symbiodinium* across the Great Barrier Reef. PLoS ONE, 8(7), e68533. doi: 10.1371/journal.pone.0068533

Van den Hoek, C., Breeman, A. M., Bak, R. P. M., & Van Buurt, G. (1978). The distribution of algae, corals and gorgonians in relation to depth, light attenuation, water movement and grazing pressure in the fringing coral reef of Curaçao, Netherlands Antilles. Aquatic Botany, 5, 1–46. doi: 10.1016/0304-3770(78)90045-1

Warner, M. E., LaJeunesse, T. C., Robison, J. D., & Thur, R. M. (2006). The ecological distribution and comparative photobiology of symbiotic dinoflagellates from reef corals in Belize: Potential implications for coral bleaching. Limnology and Oceanography, 51(4), 1887–1897. doi: 10.4319/lo.2006.51.4.1887

Warnes, G. R., Bolker, B., Bonebakker, L., Gentleman, R., Huber, W., Liaw, A., …Venables, B. (2020). gplots: Various R programming tools for plotting data. R package version 3.1.1. Avaliable at https://CRAN.R-project.org/package=gplots

Weber, M., de Beer, D., Lott, C., Polerecky, L., Kohls, K., Abed, R. M. M., … Fabricius, K. E. (2012). Mechanisms of damage to corals exposed to sedimentation. Proceedings of the National Academy of Sciences, 109(24), E1558–E1567. doi: 10.1073/pnas.1100715109

Woodhead, A. J., Hicks, C. C., Norström, A. V., Williams, G. J., & Graham, N. A. J. (2019). Coral reef ecosystem services in the Anthropocene. Functional Ecology, 1365–2435.13331. doi: 10.1111/1365-2435.13331

Wyatt, A. S. J., Leichter, J. J., Toth, L. T., Miyajima, T., Aronson, R. B., & Nagata, T. (2020). Heat accumulation on coral reefs mitigated by internal waves. Nature Geoscience, 13(1), 28–34. doi: 10.1038/s41561-019-0486-4

Yorifuji, M., Yamashita, H., Suzuki, G., Kawasaki, T., Tsukamoto, T., Okada, W., … Harii, S. (2021). Unique environmental Symbiodiniaceae diversity at an isolated island in the northwestern Pacific. Molecular Phylogenetics and Evolution, 161, 107158. doi: 10.1016/j.ympev.2021.107158

