## Supplemental figures and tables for "Community composition of coral-associated Symbiodiniaceae is driven by fine-scale environmental gradients"

**Supplemental material**

**Table 1.** Site information

| Site | Block | Depth (m) | Sedimentation (mean g/day) | Min temp ºC | Max temp ºC |
|---|---|---|---|---|---|
| 1_2 | 1 | 1.2 | 0.06602 | 21.96300 | 29.37280 |
| 1_3 | 1 | 3 | 0.02187 | 21.85184 | 28.82504 |
| 1_4 | 1 | 3.5 | 0.01931 | 21.86328 | 28.84163 |
| 1_6 | 1 | 0.7 | 0.02287 | 21.30655 | 28.94215 |
| 1_9 | 1 | 2.6 | 0.02982 | 21.79313 | 28.81257 |
| 1_10 | 1 | 3 | 0.02928 | 21.78954 | 28.88589 |
| 2_1 | 2 | 1.2 | 0.16410 | 22.31494 | 28.85388 |
| 2_2 | 2 | 0.5 | 0.04501 | 21.84382 | 28.87942 |
| 2_3 | 2 | 0.5 | 0.02779 | 21.84382 | 29.27528 |
| 2_4 | 2 | 2.4 | 2.93056 | 22.26726 | 28.62096 |
| 2_7 | 2 | 1.5 | 0.40130 | 22.30103 | 28.72309 |
| 2_8 | 2 | 2.1 | 0.27611 | 22.25235 | 28.72309 |
| 3_2 | 3 | 1.4 | 0.09804 | 21.84382 | 28.14983 |
| 3_3 | 3 | 1 | 0.05737 | 21.95887 | 28.95064 |
| 3_4 | 3 | 1 | 0.04163 | 20.85519 | 29.28453 |
| 3_5 | 3 | 0.9 | 0.03061 | 22.05129 | 29.03197 |
| 3_6 | 3 | 0.9 | 0.03796 | 21.89967 | 28.98258 |
| 3_7 | 3 | 0.7 | 0.02244 | 22.06799 | 29.16797 |
| 4_4 | 4 | 1.4 | 0.04811 | 21.92140 | 28.94923 |
| 4_5 | 4 | 1.1 | 0.03305 | 22.06524 | 28.92978 |
| 4_6 | 4 | 1.5 | 0.34257 | 21.26972 | 29.18656 |
| 4_8 | 4 | 1.2 | 0.01573 | 21.93933 | 28.83634 |
| 4_11 | 4 | 1.4 | 0.07752 | 21.84382 | 29.22060 |
| 4_14 | 4 | 1.4 | 0.16850 | 21.84382 | 28.92601 |
| 5_1 | 5 | 1.2 | 0.21580 | 20.78666 | 29.38226 |
| 5_2 | 5 | 2.7 | 0.62898 | 21.97400 | 28.81713 |
| 5_3 | 5 | 2.4 | 0.13268 | 21.89458 | 28.79825 |
| 5_6 | 5 | 3.5 | 0.41842 | 22.02388 | 27.61192 |
| 5_7 | 5 | 3.3 | 0.38405 | 21.97676 | 28.69611 |
| 5_8 | 5 | 3.2 | 0.28655 | 21.82541 | 28.74865 |

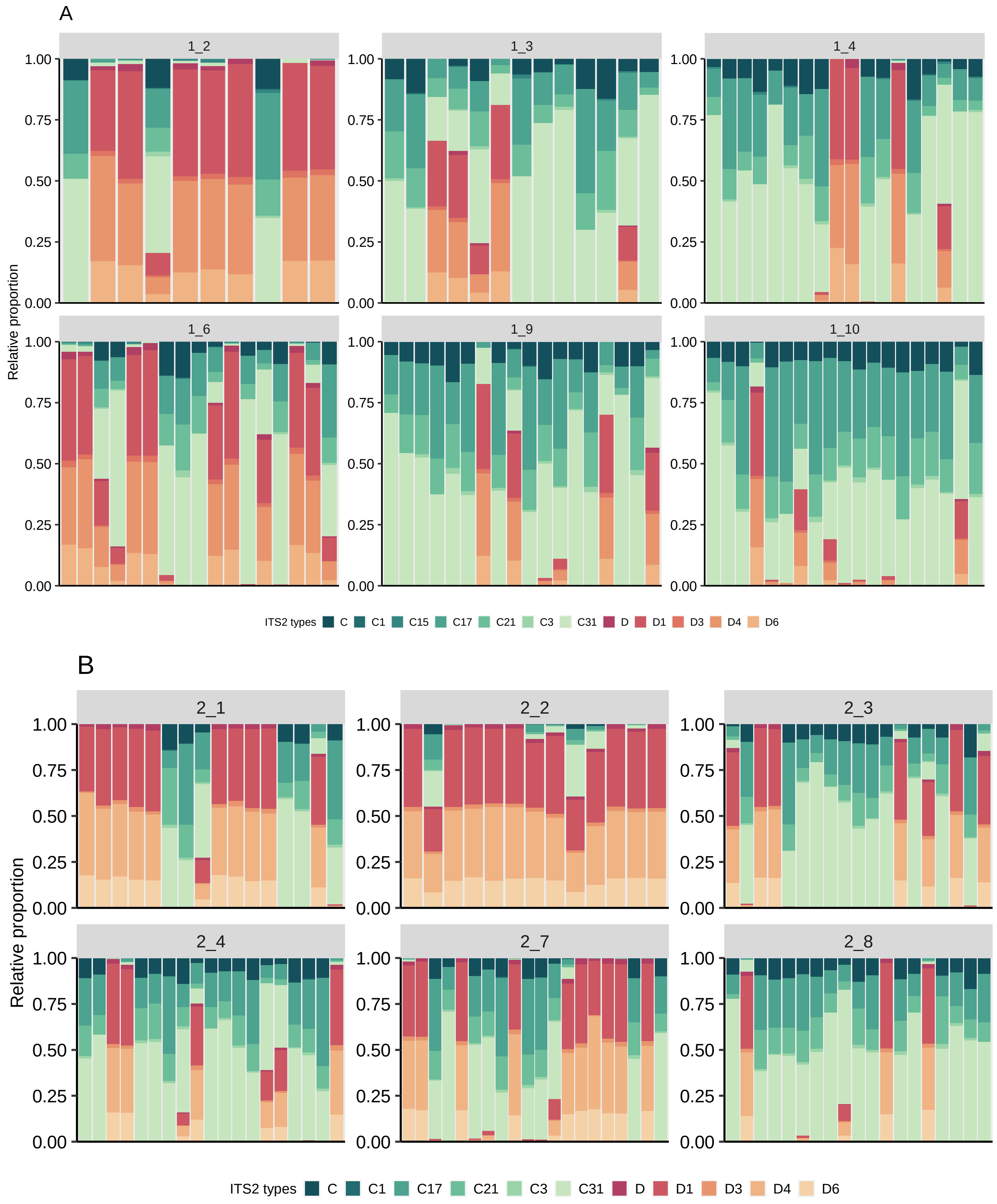
A
1_2
1_3
1_4
1_6
1_9
1_10
Relative proportion
1.00
0.75
0.50
0.25
0.00
ITS2 types C C1 C15 C17 C21 C3 C31 D D1 D3 D4 D6
B
2_1
2_2
2_3
2_4
2_7
2_8
Relative proportion
ITS2 types C C1 C17 C21 C3 C31 D D1 D3 D4 D6

C

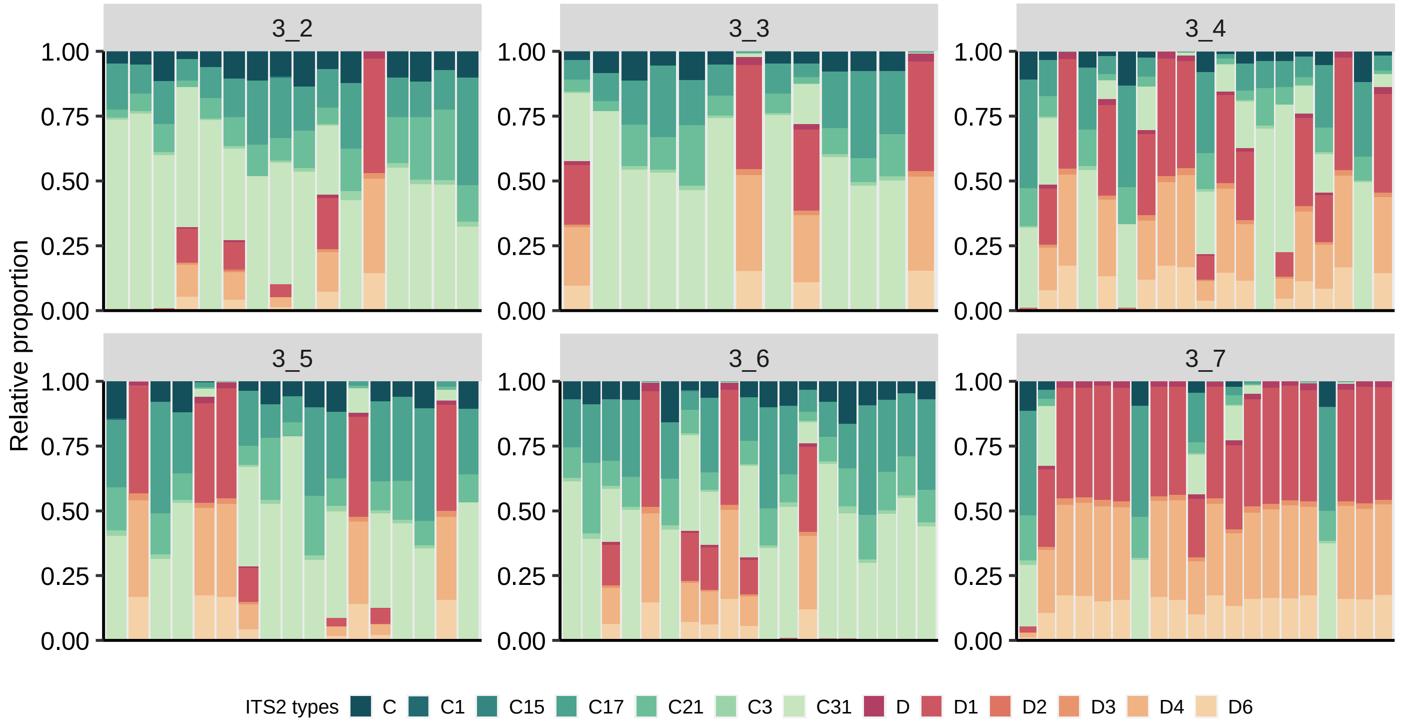
3_2
3_3
3_4
3_5
3_6
3_7
Relative proportion
0.00
0.25
0.50
0.75
1.00
ITS2 types
C
C1
C15
C17
C21
C3
C31
D
D1
D2
D3
D4
D6

D

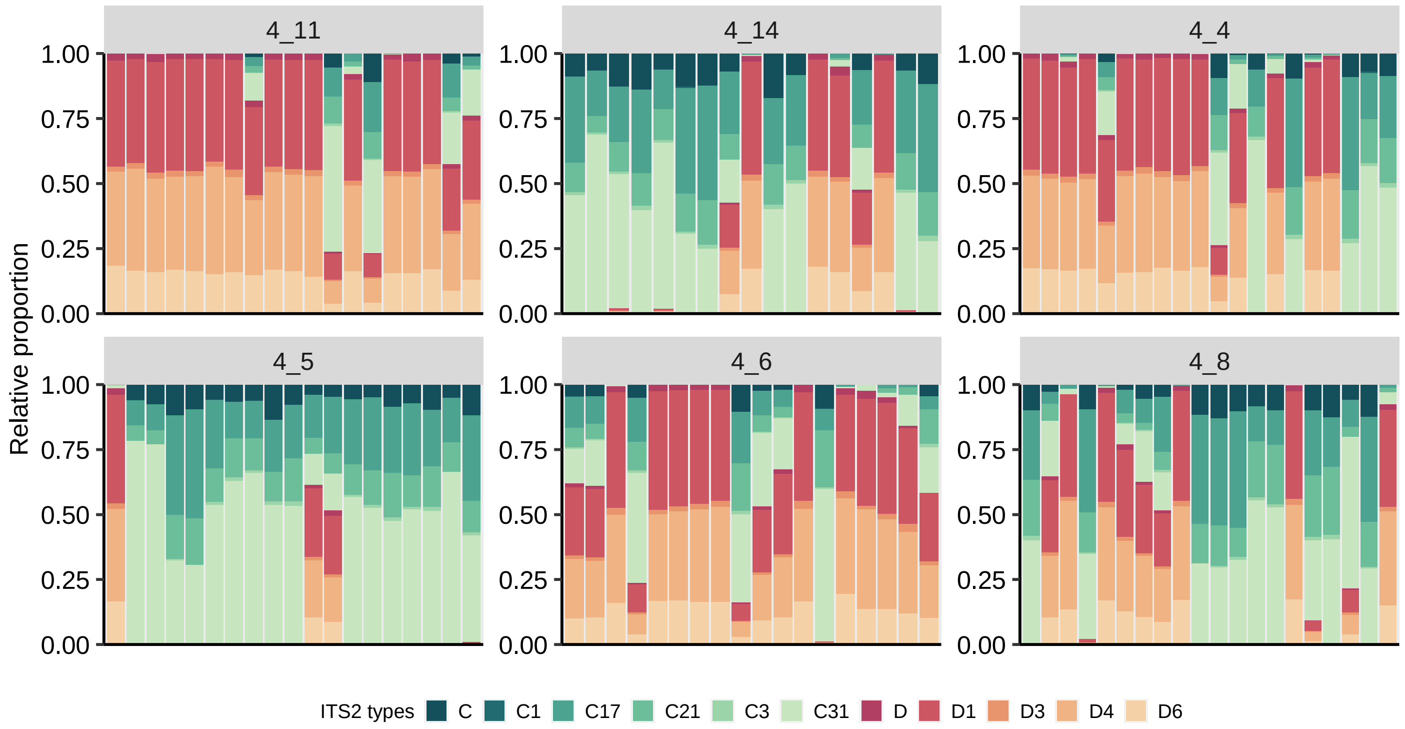
4_11
4_14
4_4
4_5
4_6
4_8
Relative proportion
0.00
0.25
0.50
0.75
1.00
ITS2 types
C
C1
C17
C21
C3
C31
D
D1
D3
D4
D6

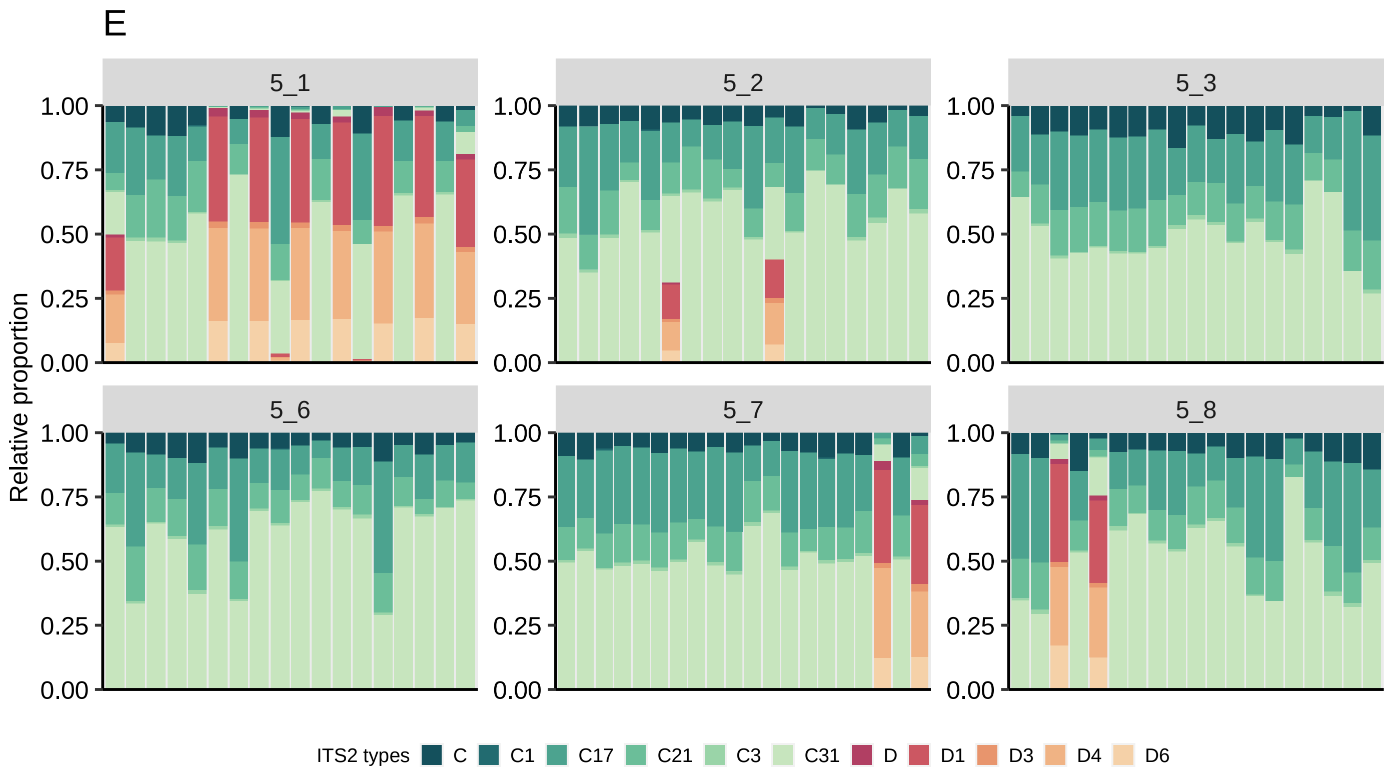


**Figure 1.** Relative proportion of Symbiodiniaceae types present in each *Montipora capitata* colony in each site. Panels shows the different blocks - block 1 (A), block 2 (B), block 3 (C), block 4 (D), block 5 (E).

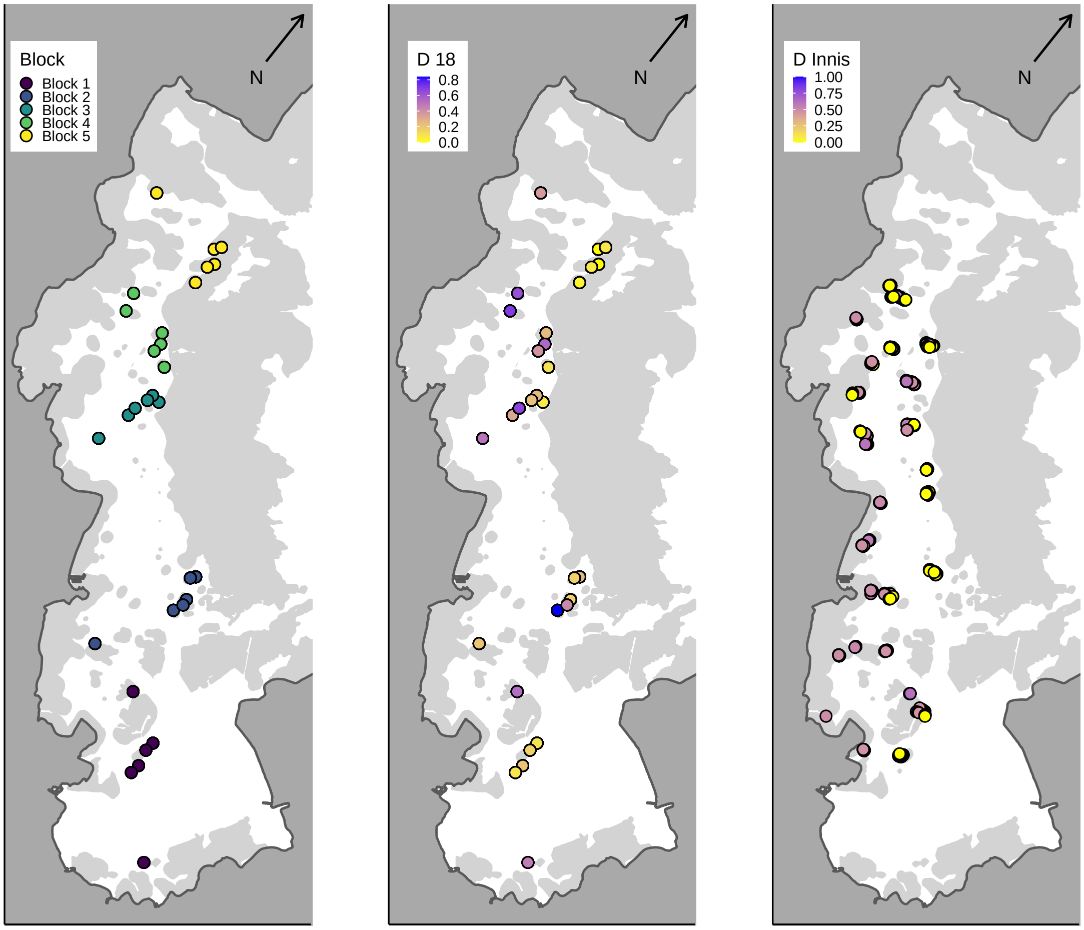


**Figure 2.** Comparison of proportion of *Durusdinium* (previous clade D). (A) Blocks used in our study, (B) proportion of *Durusdinium* in our study (C) proportion of *Durusdinium* reported in Innis et al (2018).

**Table 2.** Temperature summary statistics for each block.

| Block | Mean ºC | Maximum ºC | Minimum ºC | Daily range | Mean daily standard deviation |
|---|---|---|---|---|---|
| **1** | 25.697 | 28.946 | 21.761 | 0.244 | 0.243 |
| **2** | 25.647 | 28.834 | 22.137 | 1.246 | 0.335 |
| **3** | 25.798 | 28.928 | 21.779 | 1.871 | 0.474 |
| **4** | 25.716 | 29.008 | 21.813 | 1.261 | 0.342 |
| **5** | 25.490 | 28.675 | 21.746 | 0.746 | 0.204 |

**Table 3.** Pearson correlation coefficient of depth and temperature, and depth and sedimentation parameters used in this study.

| | Degrees of freedom | r | P |
|---|---|---|---|
| *Depth with Temperature* | | | |
| DHW | 548 | -0.4541 | <0.001* |
| Mean | 548 | -0.3393 | <0.001* |
| Max | 548 | -0.5544 | <0.001* |
| Min | 548 | 0.0311 | 0.465 |
| Daily range | 548 | -0.7425 | <0.001* |
| Mean daily standard deviation | 548 | -0.7565 | <0.001* |
| *Depth with Sedimentation* | | | |
| Mean | 548 | 0.2512 | <0.001* |
| Max | 548 | 0.2380 | <0.001* |
| Min | 548 | 0.2988 | <0.001* |
| Std | 548 | 0.2408 | <0.001* |
